# niiv: Interactive Self-supervised Neural Implicit Isotropic Volume Reconstruction

**DOI:** 10.1101/2024.09.07.611785

**Authors:** Jakob Troidl, Yiqing Liang, Johanna Beyer, Mojtaba Tavakoli, Johann Danzl, Markus Hadwiger, Hanspeter Pfister, James Tompkin

## Abstract

Three-dimensional (3D) microscopy data often is anisotropic with significantly lower resolution (up to 8×) along the *z* axis than along the *xy* axes. Computationally generating plausible isotropic resolution from anisotropic imaging data would benefit the visual analysis of large-scale volumes. This paper proposes niiv, a self-supervised method for isotropic reconstruction of 3D microscopy data that can quickly produce images at arbitrary output resolutions. The representation embeds a learned latent code within a neural field that describes the implicit higher-resolution isotropic image region. We use a novel attention-guided latent interpolation approach, which allows flexible information exchange over a local latent neighborhood. Under isotropic volume assumptions, we self-supervise this representation on low-/high-resolution lateral image pairs to reconstruct an isotropic volume from low-resolution axial images. We evaluate our method on simulated and real anisotropic electron (EM) and light microscopy (LM) data. Compared to a state-of-the-art diffusion-based method, niiv shows improved reconstruction quality (+1 dB PSNR) and is over three orders of magnitude faster (1,000×) to infer. Specifically, niiv reconstructs a 128^3^ voxel volume in 2/10th of a second, renderable at varying (continuous) high resolutions for display.

## 1 Introduction

3D imaging data is ubiquitous in scientific domains such as biology or material sciences. However, many imaging modalities like 3D electron microscopy (EM) or light microscopy (LM) have limited axial (*z*) resolution due to physical sectioning of tissue slices or optical limitations. Thus, resolution is typically much higher in the lateral directions than in the axial direction (Fig. 1a).

**Fig. 1.**
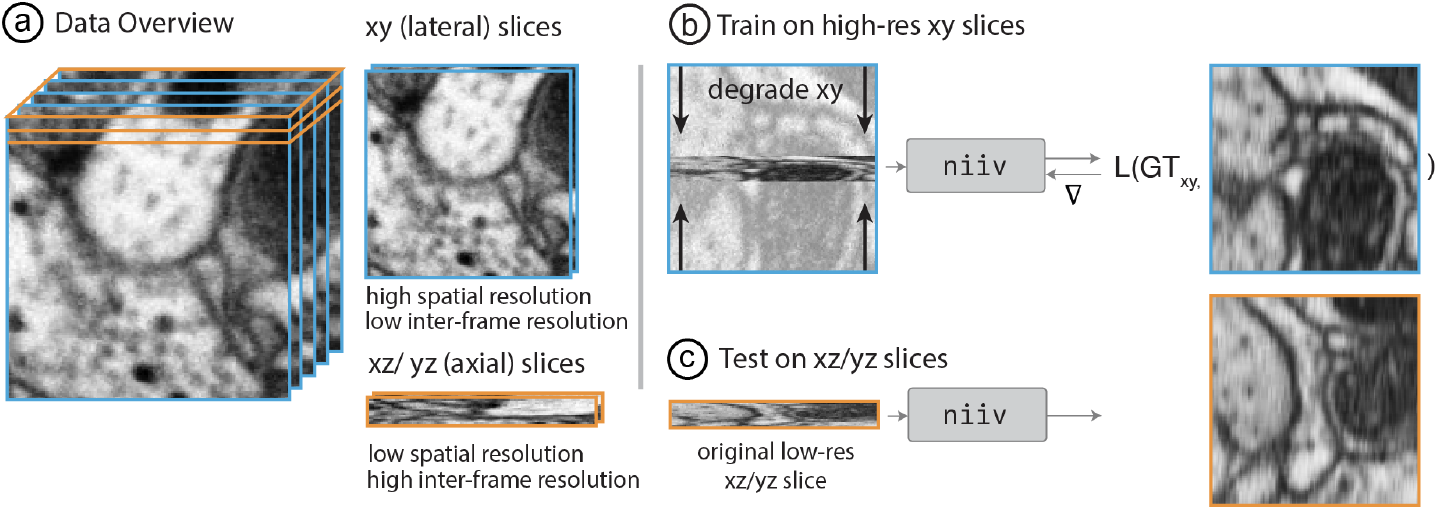
Workflow. (a) Lateral (*xy*) slices in anisotropic volumes have a high spatial but low inter-frame resolution, while axial (*xz*/*yz*) slices have a low spatial but high inter-frame resolution. (b) Thus, niiv is trained to predict high-resolution lateral slices from artificially downsampled input. (c) niiv infers isotropic volumes at test time by predicting high-resolution axial (*xz*/*yz*) slices from their low-resolution counterparts.

Downstream tasks like interactive visual analysis would benefit from high-resolution isotropic volumes, but these can be extremely costly or impossible to obtain. Several computational methods attempt to generate isotropically resolved volumes [31,30,11,21,34,35,8,5]. Some approaches [31,30] require specific shape priors, such as exact point spread functions (PSFs), which are difficult to measure in practice. Machine learning approaches like *diffusion* can model complex data distributions but require copious training data and are slow to infer [11,21]—requiring multiple minutes or even hours to reconstruct a small isotropic volume.

At the same time, anisotropic imaging volumes grow in size every year, containing terabytes [36,27] or even petabytes [24] of imaging data. Reconstructing isotropic volumes is typically an offline postprocess after image acquisition, but with increasing data sizes this becomes infeasible. For instance, Lee et al. [12] take up to three minutes to reconstruct a 128^3^ voxel volume. Instead, isotropic volumes should be reconstructed on-demand and locally at interactive rates from anisotropic data for visual inspection. Faster neural implicit reconstruction approaches are either supervised [5] or rely on bilinear latent interpolation [34] with fixed averaging, thus limiting the field of view per queried latent.

Thus, we propose an interactive self-supervised method to reconstruct isotropic volumes from anisotropic data, which also enables learnt latent interpolation using a local attention mechanism. Building on recent advances in neural field representations [32,10,19,25], our model uses a super-resolution encoder [14] to relate a low-resolution axial slice to a plausible high-resolution image via a latent space [2,3]. A multi-layer perceptron (MLP) decodes a set of latent codes into a high-resolution axial slice sample, with local attention-guided latent interpolation creating an output image at any pixel resolution (Fig. 2). Both encoder and decoder are trained end-to-end on simulated anisotropic slices by downsampling isotropic lateral slices (Fig 1b). Inference is fast, requiring only 2/10th of a second to generate a 128^3^ voxel volume, allowing interactive isotropic visual inspection.

**Fig. 2.**
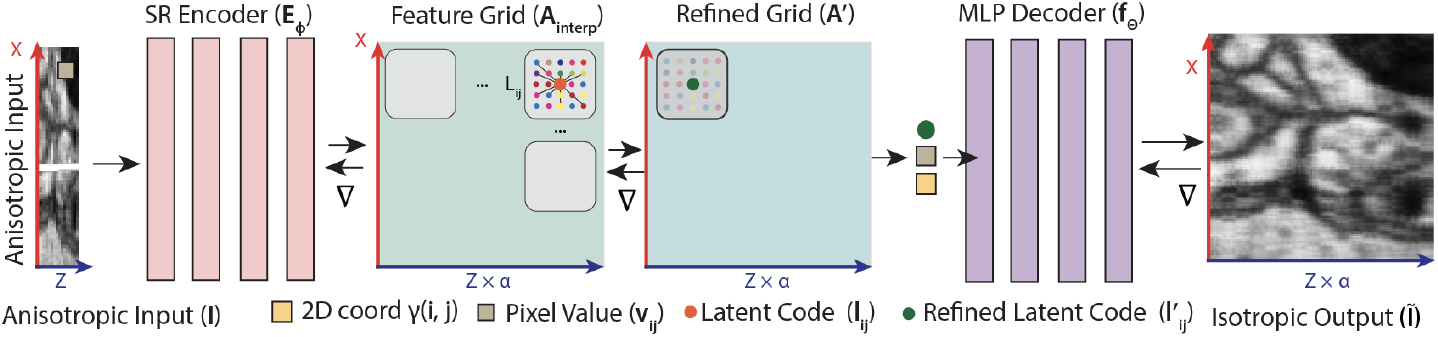
Model overview. A super-resolution encoder *E*_*ϕ*_ [14] embeds the anisotropic input slice into latent codes placed within a 2D spatially-referenced grid *A. A* is bilinearly interpolated (*A*_interp_) at output coordinates and subsequently refined using local attention (*A*^*′*^). We predict a pixel value in the output by querying *A*^*′*^ at its coordinate (*i, j*) and concatenating the resulting latent code *l*_*i*_^*′*^_*j*_ (green) with the positionally encoded coordinate (*i, j*) (yellow) and the value at (*i, j*) in the input (beige). The decoding function *f*_*θ*_, parametrized by an MLP, predicts the respective pixel value.

For validation, we compare niiv against bilinear upsampling and three current self-supervised methods, including two neural-field [25,34], and one diffusion-based approach [12]. Our approach shows quality improvements over all and quantitative improvements by +1 dB over the diffusion model. We demonstrate this through peak-signal-to-noise ratio (PSNR) computations in a sweep of a frequency-clipped Fourier domain, which offers a more robust metric than pixelwise PSNR to noise in the training data. For computational efficiency, niiv is 1,000× faster than the diffusion model, 666× faster than the SIREN baseline considering volume-specific pertaining times and achieves similar inference speeds to Zhang et al. [34].

## 2 Related Work

### Isotropic Volume Reconstruction

Recent self-supervised approaches [21] [12,15,13] train 2D diffusion models to learn the distribution of high-resolution lateral images. During reconstruction, they use low-resolution axial slices as priors for the backward diffusion process to predict missing volume information. While diffusion models achieve high-quality results, their usability is limited by compute-intensive training and time-consuming inference. In contrast, our approach improves reconstruction quality while inferencing three orders of magnitude faster than the diffusion baseline [12]. Recently, Zhang et al. [34] also use neural representations for anisotropic volume reconstruction with bilinear latent interpolation. Our approach additionally enables learning latent interpolation through a local attention mechanism. Deng et al. [4] learns a degradation model that generates realistic low-resolution, high-resolution training pairs. These pairs are used to train a reconstruction model like Iso-Net-2 [31]. Both models are trained independently, leading to complex training setups. On the other hand, supervised approaches [8,5] show high-quality results but are challenging to use in practice since isotropic volumes are required at training time. Other methods use video transformers [7], optical flow field interpolation [1], or standard ConvNets [31,35], which are limited to a specific output pixel resolution, whereas niiv can be decoded at any resolution.

### Neural Implicit Super-Resolution

Encoding spatial information through implicit neural representations (INR) [20,32] has proven to be useful in areas, such as inverse graphics [19,25], shape representation [26,33,9], video encoding [10], and super resolution [2,3,18,34,35,5]. Our approach builds upon local implicit image functions (LIIF) [2], which allows the sampling of images at arbitrary resolution while retaining high-quality visual details. Critically, our approach differs from related approaches [35,34] by using attention guided latent refinement mechanism that enables learnt feature interpolation over a local latent neighborhood (Fig. 2).

## 3 Methodology

### Problem Statement

Given a sampling of a volume *V* with isotropic *xy* axes and anisotropic *z* axis, we aim to learn a model *g* that reconstructs an isotropic *z* sampling of a volume 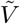 purely by self-supervision. We assume that the volume sampling contains a physical medium whose distribution of material can be effectively modeled by observing local statistics of the *xy* samplings. Given the low-resolution anisotropic slice 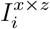 as input, *g* must reconstruct a plausible high-resolution isotropic *xz* slice 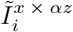, where *i* denotes a slice from the input volume sampling and *α* is the axial anisotropy factor (e.g., 8). Then, 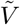 is constructed by stacking predicted high-resolution slices 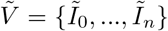. The same approach applies equally to both *xz* and *yz* slices.

### INR with Learned Latent Interpolation

Our hybrid neural field [32] uses latent codes within a 2D space to encode the high-resolution slice. First, a convolutional super-resolution encoder *E*_*ϕ*_ with parameters *ϕ* embeds a low-resolution axial slice *I* ∈ ℝ^*x×z*^ into a 2D referenced *d*-dimensional latent grid *A* ∈ ℝ^*x×z×d*^, of latent codes *l*. Next, we bilinearly interpolate *A* at the desired output coordinates, such that *A*_interp_ ∈ ℝ^*x×*(*zα*)*×d*^. Next, we locally refine *A*_interp_ by attending each latent *l*_*ij*_ ∈ *A*_interp_ to its local neighborhood *L*_*ij*_ (Fig. 2). We name refined latent codes 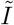, and the refined latent grid *A*^*′*^.

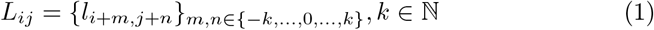

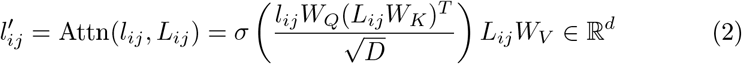

*W*_*Q*_, *W*_*K*_, *W*_*V*_ and *D* define the attention mechanism’s trainable weight matrices and dimensionality, respectively [29]. *σ* represents the softmax function. Finally, a MLP decoder *f*_*θ*_ with parameters *θ* (Fig. 2) produces reconstructed image intensities *Ĩ* at an output pixel coordinate (*x, y*):

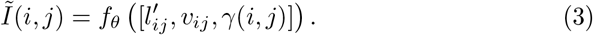

(*x, y*) is encoded using a 2-band frequency basis *γ*(*i, j*), and *v*_*ij*_ is the pixel value obtained by bilinearly interpolating *I* at (*i, j*). Thus, niiv can adapt flexibly to the requirements of interactive display across devices, unlike other approaches [8,31]. *E*_*ϕ*_ and *f*_*θ*_ are shared between volumes in the training and test dataset. We use the mean absolute error loss function (MAE) during training.

### Simulating axial degradation

During training, we use artificially degraded *xy* images 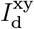 as model inputs and supervise outputs with the respective high-resolution *xy* slices 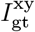. We apply a function *d* that aims to simulate degradation along the *z* axis such that 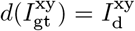. Here, we define *d* as an average pooling operator. In principle, other degradation models [4,12,6] can be applied based on the specific application and imaging domain.

## 4 Evaluating Reconstructions with Noisy Ground Truth

Evaluating niiv brings challenges, as only noisy ground truth exists, leading to uninformative PSNR values (Fig. 3a). Thus, a method that perfectly reconstructs lateral slices is overfitting to the noise; this is especially problematic in low-data regimes with powerful data-fitting models [12]. We wish to assess the reconstruction of biological structures despite the noise. Prior research has addressed this problem by downscaling the data to diminish noise [8] at the cost of sacrificing resolution. We propose calculating the PSNR in the Fourier domain where it is easier to separate high-frequency components of signals [6] such as noise (Fig. 3b), where we vary a cutoff frequency *f*_*cutoff*_ (Fig. 3c) across a range of values to observe the quality across frequencies. For example, if a method only achieves a greater PSNR than another at high *f*_*cutoff*_ but not at low *f*_*cutoff*_, then it is likely overfitting the noise. Given Parseval’s theorem [22] and the unitary nature of the Fourier transform ℱ, we can directly compute the PSNR in the frequency domain, sidestepping the inverse transformation to the spatial domain.

**Fig. 3.**
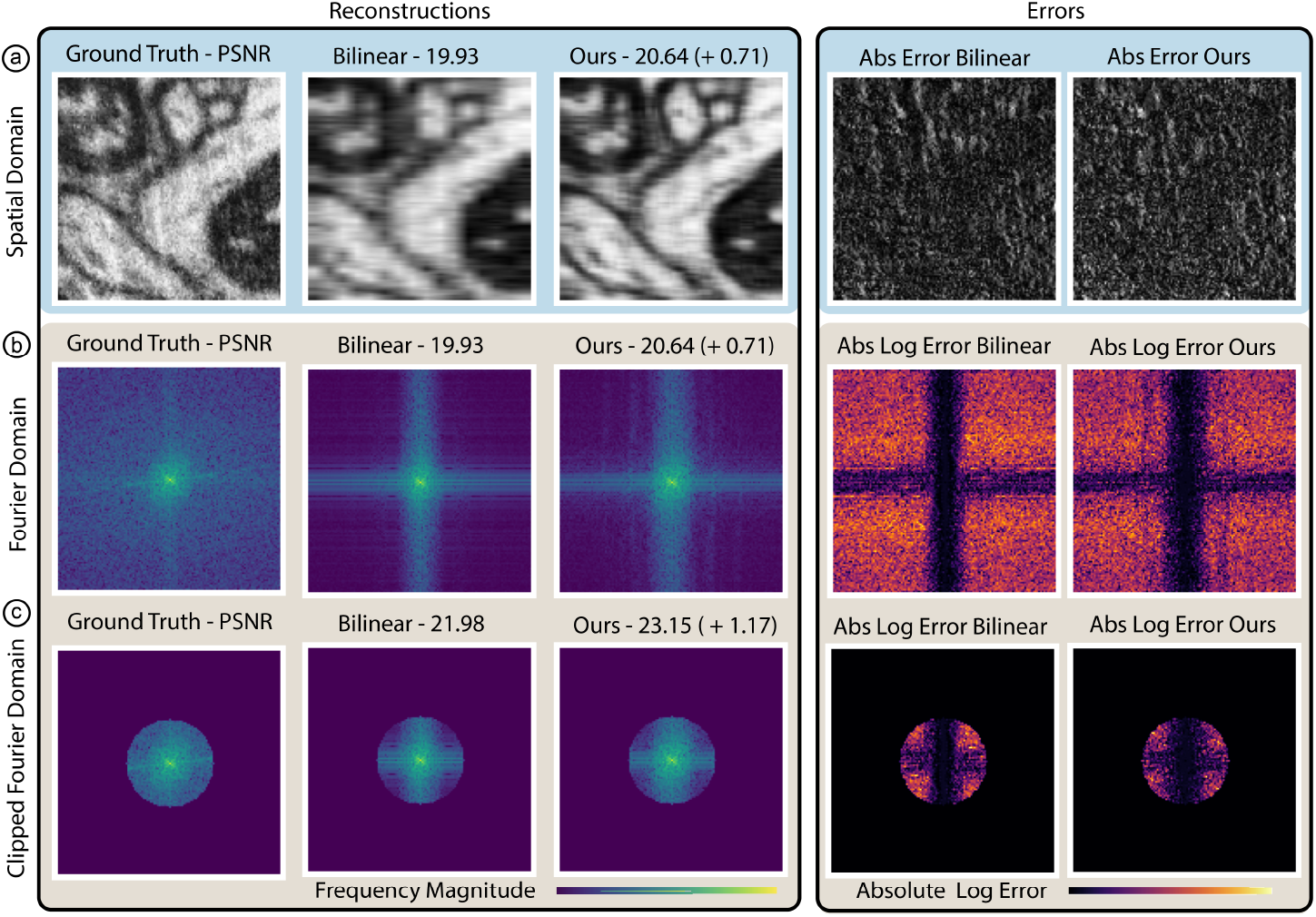
Fourier space separates incorrect from correctly reconstructed frequencies. (a) In the spatial domain, reconstruction errors are randomly distributed for the baseline and our reconstruction. (b) Visualizing errors in the Fourier domain (right column) separates erroneous (bright) and correctly reconstructed frequencies (dark). (c) By cropping the Fourier domain’s high frequencies, the PSNR is not perturbed by frequencies corresponding to noise, yielding more informative PSNR values.

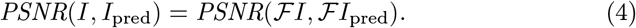

We now incorporate the clipping operation in the Fourier domain, denoted by a low-pass operator *L* that discards frequencies above *f*_*cutoff*_. Given the above relationship, we receive

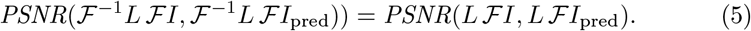

That is, we can transform both images into the Fourier domain, apply the clipping operation via *L*, and then compute the PSNR directly in Fourier space.

## 5 Experiments

### 5.1 Data and Implementation Details

We demonstrate the effectiveness of our approach on the publicly available FlyEM Hemibrain [23], FAFB [36] EM datasets and also ablate against LM approaches like LICONN [28] (Fig 5b). While the Hemibrain contains the central brain region of Drosophila melanogaster imaged at isotropic 8 × 8 × 8 nm pixel resolution, we downsample the data to 8× anisotropy along the *z* -axis through average pooling. FAFB shows the entire brain of a female adult fruit fly at naturally 5× anisotropic 8 × 8 × 40 nm pixel resolution. We randomly sample 400 subvolumes (128^3^ pixels in the Hemibrain and 130^3^ pixels in FAFB) and separate them into training (*N* = 350) and test datasets (*N* = 50). All metrics are reported on entire volumes rather than individual images. Our method is implemented in PyTorch, and all tests were performed on a single NVIDIA RTX 3090 Ti GPU. All experiments use the EDSR [14] super-resolution encoder without upsampling modules, six residual blocks, and 64-dimensional output features. The MLP is five layers deep, each 256 neurons wide. We train our model for 900 epochs using the Adam optimizer and a learning rate of 5 × 10^*−*5^.

### 5.2 Qualitative and Quantitative Comparison

To showcase our method’s suitability for interactive reconstruction, we capped the GPU memory usage at 4 GB for all methods, reflecting a mid-tier laptop’s capacity. Within this constraint, our approach significantly outperforms the diffusion baseline, delivering inference speeds up to three orders of magnitude faster (0.225 vs. 264 seconds) for an anisotropy reconstruction task with anisotropy *α* = 8 on 128^3^ volumes (Hemibrain). The advantage is due to Diffusion-EM’s slow iterative inference process and the need to enforce frame-by-frame consistency for the probabilistic reconstruction process by conditioning each slice inference on a latent code retrieved from the previous slice, prohibiting batch processing. We also outperform SIREN [25] as it requires separate pretraining for each subvolume, leading to costly inference on unseen data (Table 1). Additionally, we achieve better qualitative results (CF PSNR, PSNR, SSIM) than Zhang et al. [34], while being slightly slower due to the additional computations required to compute attention-based latent interpolation. Comparing reconstruction quality, in contrast to the baselines, our model can reconstruct fine details (Fig. 4a) with sharp edges (Fig. 4b). While the diffusion results visually look sharp, small details are often reconstructed incorrectly, explaining the lower metric scores. Diffusion EM also fails for volume sizes, not in {2^*i*^} (Fig. 4b).

**Table 1.** Hemibrain Evaluation. We report clipped Fourier PSNR (CF PSNR) with *f*_*cutoff*_ = 25, regular PSNR, and SSIM. We differentiate between volume-specific pretraining time and inference time (Pre/Infer). The highest scores are highlighted in red.

| Hemibrain Data | CF PSNR ↑ | PSNR ↑ | SSIM ↑ | Time (Pre/Infer) ↓ | VRAM (GB) |
| --- | --- | --- | --- | --- | --- |
| Nearest | 23.16 | 20.42 | 0.50 | — | — |
| Bilinear | 23.53 | 20.92 | 0.51 | — | — |
| SIREN [25] | 19.55 | 18.24 | 0.34 | (150s/0.003s) | 3.9 |
| Diffusion EM [12] | 23.24 | 20.65 | 0.56 | (-/264s) | 3.9 |
| Zhang et al. [34] | 24.86 | 21.63 | 0.57 | (-/0.113s) | 3.4 |
| Ours | 25.06 | 21.72 | 0.57 | (-/0.225s) | 3.9 |

**Fig. 4.**
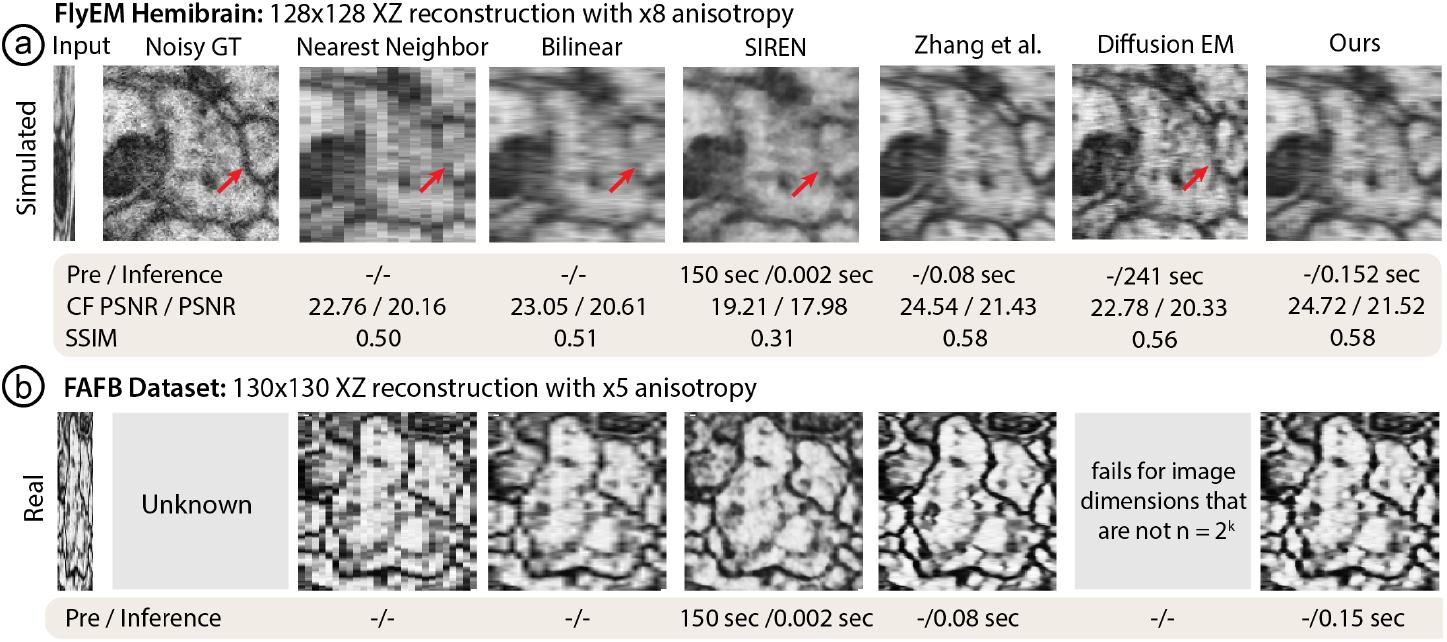
Qualitative comparison of simulated (a) and real (b) anisotropic data. We achieve up to three orders of magnitude faster inference compared to diffusion baselines [12] and other neural implicit approaches like SIREN [25] while also reconstructing smaller structures with higher fidelity (see arrows).

### 5.3 Ablation Studies

#### Fourier PSNR

We tested the effect of the Fourier clipping threshold on the PSNR (Fig. 5a). If no clipping threshold is applied, the PSNR of our method and the baselines are low and close together due to the random image noise in the ground truth data. However, the black box (Fig. 5a) indicates a clipping window in Fourier space where image quality differences for the SIREN, Diff-EM, and Bilinear baselines are more accurately represented through the PSNR.

#### Data Modality

We use a recent, near isotropic voxel-size (9.7 × 9.7 × 13 nm) expansion LM dataset [28] of a mammalian hippocampus and simulate 8× anistropy using average pooling as a degradation model (Fig. 5b). Next, we train on 350 randomly sampled volumes and reconstruct 50 unseen 128^3^ voxel test volumes at isotropic voxel size. Fig. 5b shows input and GT images and also compares our results with nearest- and bilinear interpolation. We find that niiv produces sharper images compared to bilinear interpolation also for LM data.

**Fig. 5.**
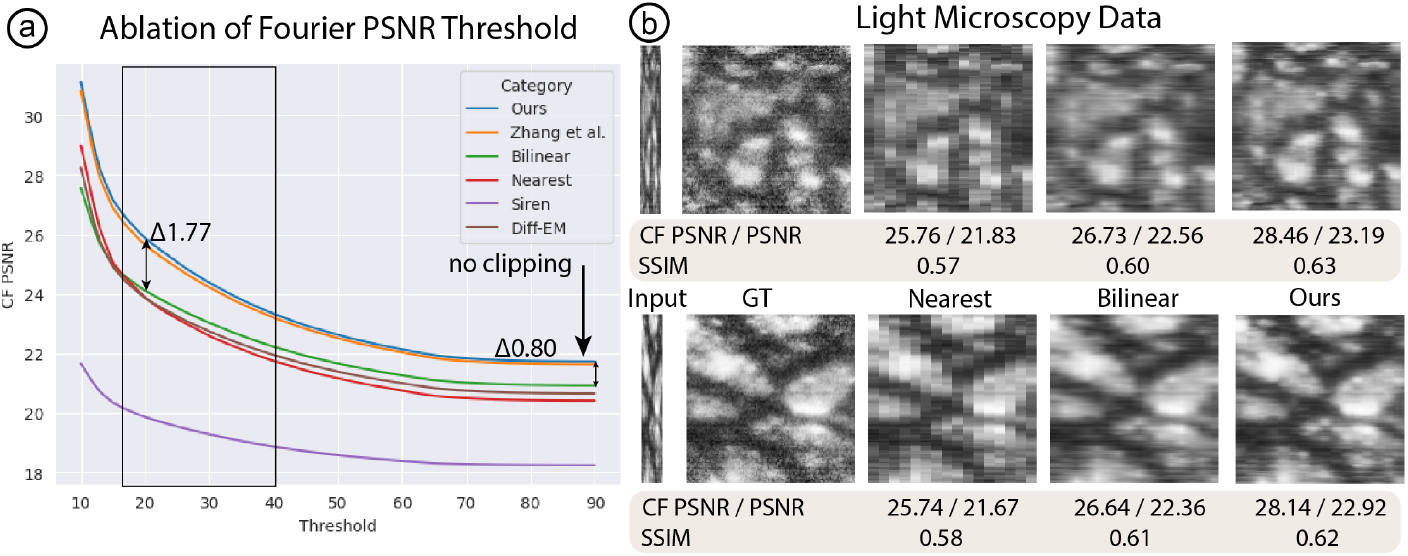
Ablation studies. (a) While the unclipped PSNR in the Fourier domain shows no significant difference between our approach and the baselines (*Δ*0.6), the difference becomes more evident for thresholds *f*_*cutoff*_ ∈ [14, 40] (*Δ*1.63). (b) We also test niiv on recent expansion light microscopy data [28] with simulated 8× anisotropy. *f*_*cutoff*_ = 25.

## 6 Conclusions and Future Work

Interactive isotropic rendering of anisotropic data is useful for large-scale data visual inspection tasks. Thus, we demonstrate that neural fields and encoder-based superresolution representations are promising for fast and flexible self-supervised volume reconstruction. We propose two avenues for future work. First, integrating machine-learning elements like our approach into low-power Web-based image-rendering tools such as neuroglancer [16] or Viv [17] would rapidly deploy these advances. Second, future work should investigate if latent image representations express higher-level semantically-interpretable morphological features, as these could be useful in downstream tasks such as tissue classification.

