## Supplemental Text for "niiv: Interactive Self-supervised Neural Implicit Isotropic Volume Reconstruction"

### 1 Details on Fourier PSNR

Parseval's theorem states that, given proper normalization, the $L_2$ norm of a signal in the spatial domain is the same as the $L_2$ norm of the signal in the frequency domain. For a discrete 1D function with $N$ samples,

$$\sum_{n=0}^{N-1} |x_n|^2 = \frac{1}{N} \sum_{n=0}^{N-1} |X_n|^2 . \tag{1}$$

Here, the discrete coefficients of the transform are given by

$$X_k = \sum_{n=0}^{N-1} x_n e^{-i2\pi \frac{nk}{N}} , \tag{2}$$

$$x_k = \frac{1}{N} \sum_{n=0}^{N-1} X_n e^{i2\pi \frac{nk}{N}} . \tag{3}$$

Here, $x$ and $X$ are discrete complex functions with $N$ samples that form a Fourier transform pair, i.e., we have $X = \mathcal{F}x$ and $x = \mathcal{F}^{-1}X$, where $\mathcal{F}$ denotes the Fourier transform. We note that here, as is often customary, the unitary normalization factors of $1/\sqrt{N}$ in both $X_k$ and $x_k$ have been substituted by a single (non-unitary) normalization factor of $1/N$ in $x_k$. We will compensate for this factor in all PSNR computations to obtain the same result as that of the unitary transform. For correctly incorporating the normalization factors 1 and $1/N$ used in Eqs. 2 and 3, respectively, we note that

$$\log_{10}\left(\frac{1}{N} \cdot MSE\right) = \log_{10} MSE - \log_{10} N. \tag{4}$$

As the MSE already normalizes by a factor of $1/N$, we compute the PSNR between the two images $A := L\mathcal{F}I$ and $B := L\mathcal{F}I_{\text{pred}}$ in the Fourier domain by coefficients $A_n$ and $B_n$ as

$$PSNR(A, B) = 20 \log_{10} (I_{max} \cdot N) - 10 \log_{10} \sum_{n=0}^{N-1} |A_n - B_n|^2 . \tag{5}$$
